# Lower-Limb Mechanical Power Accounts for Running Energy Expenditure and Enables Single-IMU Estimation

**DOI:** 10.64898/2026.07.08.737376

**Authors:** Jinsung Jung, Hyerim Lim, Sukyung Park

## Abstract

Energy expenditure (EE) during running depends on the interplay between active muscle work and elastic energy storage and return, yet the relative contribution of mechanical power to EE remains debated. Quantifying the relative contributions of segment-level mechanical power can provide a way to address this debate. In this study, we aimed to quantify how segment-level mechanical power contributes to EE during running and to demonstrate that these mechanistic insights support wearable-based EE estimation. Joint dynamics and respiratory gas-based EE were collected from healthy young adults running at multiple speeds. Scale factors were derived to quantitatively link efficiency-weighted segment power to measured EE. The stance leg consistently showed the strongest correlation with EE, and this dominance was preserved across speeds. Including swing-leg hip power further improved accuracy. Scale factors were approximately 0.45, suggesting that active muscle work and elastic energy return contribute comparably to the mechanical power associated with EE. Using a lightweight machine learning model, stance-leg and swing-leg hip joint power were reconstructed from a single sacral IMU, enabling accurate EE prediction. These findings demonstrate that lower-limb mechanical power is a robust predictor of running EE, supporting both the extensibility of biomechanically-informed frameworks and wearable-based EE monitoring.

## Introduction

Energy expenditure (EE) during running reflects the metabolic cost of muscle activation required to meet the mechanical demands of locomotion, including joint mechanical power, elastic energy interactions, and collision related losses. Understanding these mechanisms is important for evaluating locomotor efficiency, monitoring physical activity, or informing movement strategy selection^1–3^. However, how these mechanical demands translate into metabolic cost, particularly the extent to which lower-limb mechanical power accounts for EE, remains debated. Mechanical power refers to the net power performed at the external or joint level as computed from inverse dynamics, and therefore represents the purely mechanical output of the system without directly accounting for isometric force generation or elastic energy recoil. In contrast, EE represents the total metabolic cost required to generate muscular force and compensates for dissipative losses. For example, isometric muscle force production incurs metabolic cost even though it does not produce mechanical power^4^. Conversely, joint motion produced by muscle contraction both generates mechanical power and consumes metabolic energy^5^, whereas elastic tissues can generate mechanical power without metabolic cost^6^. A long-standing view is that the metabolic cost of running is primarily determined by quasi-isometric muscle action required to support body weight, with joint mechanical power contributing only modestly^7^. Consistent with this perspective, EE has often been examined in relation to speed-related parameters such as running velocity^8^, ground contact time^7^, and step frequency^9^. In contrast, other studies have highlighted associations between joint kinetics and EE, including the mechanical demands of swinging the lower-limb segments^10^ and the production of positive and negative power at specific joints^11–13^. More recently, whole-body dynamics analyses have suggested that mechanical power generated by active muscle may account for more than half of total EE even when elastic recoil is considered^14,15^, renewing interest in the role of mechanical power in running energetics.

However, quantitative evidence regarding how individual body segments contribute to EE within whole-body mechanical power, that is, the relative contribution of segment-level mechanical power from the torso, upper limbs, and lower limbs to EE, remains limited. In walking, sagittal plane stance-leg joint power has been shown to be directly proportional to EE, and this relationship can be quantified through a proportional constant referred to as a scale factor^16^. In contrast, running kinetics differs qualitatively from walking due to its shorter stance duration, greater collision losses, the presence of a flight phase, and rapid changes in torque rate around the swing-leg hip, all of which lead to distinct patterns of joint power generation and dissipation. Running physiology also differs from walking. Elastic energy storage and return by tendons plays a prominent role^17^, providing a means of generating mechanical power without metabolic cost^18,19^. In addition, substantial energy is dissipated by soft tissues during ground collision, and the body must offset this loss to maintain steady-state running, which requires additional metabolic cost^20,21^. Given these characteristics, it remains uncertain whether the dominant contribution of stance-leg mechanical power observed in walking persists during running, a question that requires direct investigation.

If the relationship between segment-level mechanical power and EE can be established, such relationships may serve as low-dimensional indicators that represent metabolic cost. This, in turn, could enable practical EE monitoring in real world settings. Indeed, in our previous study on walking, we demonstrated that the relationship between segment-level mechanical power and EE can be leveraged to enable practical EE estimation using a single sacral IMU sensor^16^. In contrast to conventional data-driven approaches, which often exhibit systematic underestimation at high EE levels and sensitivity to signal noise^22–25^, that study provided direct evidence for the effectiveness of a biomechanically-informed approach for estimating physiological signals such as EE.

In this study, we investigate the extent to which segment-level mechanical power explains EE during running. Specifically, we address three key questions. (i) Does the dominant contribution of stance leg to EE observed in walking persist in running, despite the qualitatively different mechanics of the two gait modes? (ii) Are there additional segments that significantly improve explanatory value beyond the stance leg? (iii) How do the scale factors derived in the estimation process differ from those observed in walking, and what biomechanical meaning does these differences carry in terms of energetics? To answer these questions, we collected joint kinematics, joint kinetics, and respiratory gas–based EE from healthy young adults across multiple running speeds. We first examined the relative contributions of segment-level mechanical power of the stance leg and the swing leg to whole-body mechanical power. Scale factors linking muscle efficiency–weighted mechanical power to EE were then derived, providing quantitative insight into the contribution of active muscle work to metabolic cost. Building on these mechanistic relationships, we validated whether the previously proposed single IMU-based biomechanically-informed EE estimation framework can be extended to running, thereby assessing both the generality of the relationship and its practical applicability.

## Methods

This study consisted of two parts: experimental data collection and analytical modeling. The experimental data were collected to characterize the relationship between mechanical power and measured EE across a range of running speeds. The analytical modeling included both the derivation of segment-specific scale factors from motion capture data and the estimation of EE from a single IMU using a machine learning framework.

### Experimental data collection

Experimental data were collected to analyze segmental and planar contributions to whole-body mechanical power and to develop and validate the IMU-based EE estimation framework (Fig. 1*a*). All experimental procedures were approved by the Institutional Review Board (KH2023-250), and written informed consent was obtained from all participants prior to testing. All methods were performed in accordance with relevant ethical guidelines and regulations. Eight healthy adult males participated (age: 24.3 *±* 4.4 years; height: 174.8 *±* 5.8 cm; mass: 69.3 *±* 8.8 kg). None of the participants had a history of lower-limb musculoskeletal disorders. All participants were recreational runners who ran at least 25 km per week and could complete a 10 km run within 55 minutes. Participants were recruited from the local community. Participants ran on an instrumented treadmill at five randomized speeds (2.5, 2.8, 3.1, 3.4, and 3.7 m/s). Each trial lasted 8 minutes, and only the final 3 minutes were used for analysis to ensure steady-state metabolic conditions. The respiratory exchange ratio (RER) was continuously monitored to ensure that it remained below 1 throughout the running trials. The mean RER was 0.91 *±* 0.07, confirming that substantial anaerobic contribution did not occur. Participants were provided 10–30 minutes of rest between trials, during which heart rate, VO2, and VCO2 were monitored to confirm full recovery before proceeding. Whole-body kinematics were captured at 100 Hz using a 13-camera motion capture system (MX-T50, VICON, UK) with 40 reflective markers placed on anatomical landmarks. Ground reaction forces were recorded at 1000 Hz using an instrumented treadmill (FP6012, Bertec, USA). A single sacrum-mounted IMU (Opal, APDM, USA) recorded triaxial accelerations and angular velocities at 100 Hz. All motion capture, force plate, and IMU signals were low-pass filtered using a zero-phase 4th-order Butterworth filter with a cutoff frequency of 10 Hz, consistent with prior work on IMU-based joint dynamics estimation. Metabolic energy expenditure was obtained using indirect calorimetry (K5, Cosmed, Italy). EE was calculated from respiratory gas exchange using standard equations^26^, applying a fixed conversion factor relating O2 consumption and CO2 production to EE. Net EE was defined as the difference between average EE during the final 3 minutes of running and average resting EE measured during quiet standing. This net EE was used as the measured EE in all analyses.

**Figure 1.**
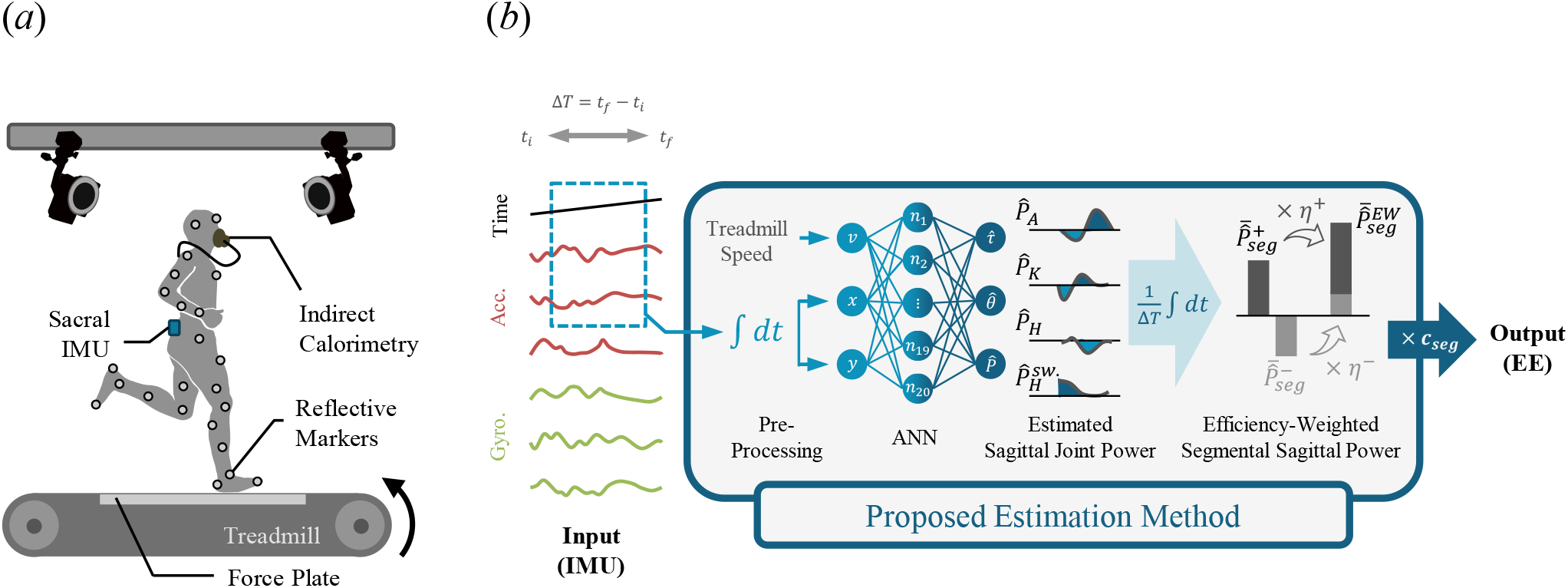
(*a*) Experiment setup. The whole-body motion capture system, a force plate, a sacrum-mounted IMU, indirect calorimetry were used. (*b*) Schematic of the proposed EE estimation method. The proposed method uses IMU data to estimate the segmental sagittal joint power as an intermediate biomechanical variable, which is then efficiency-weighted and regression-scaled to compute the whole-body EE.

### Analysis and modeling

The analytical framework consisted of two main components. The first component involved deriving segment-specific scale factors by quantifying efficiency-weighted mechanical power and regressing these values against measured EE. The second component involved estimating EE from single IMU data using a machine learning model that reconstructs stance-phase joint dynamics as intermediate variables. The procedures follow the methodology established in our previous work^16^, and the relevant steps are summarized here. Walking data from this prior dataset were also used for comparison with running, specifically to evaluate segmental and planar contributions to whole-body joint dynamics and to compare swing-leg joint power patterns. These data were processed using the same joint dynamics workflow (inverse dynamics, joint power computation, and efficiency-weighted mechanical power), and only the computed joint-level mechanical power quantities were used in the present analysis.

#### Scale factor derivation from efficiency-weighted joint power

Joint torques (*τ* _*joint*_), joint angular velocities 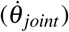, and joint mechanical powers (*P*_*joint*_) were computed from synchronized motion capture and ground reaction force data using Visual3D v6 (HAS-Motion, Canada). Following established procedures, positive and negative joint powers were time-integrated to obtain time-averaged mechanical power, and muscle efficiency-weighted joint power was computed as follows,

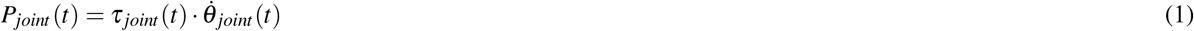

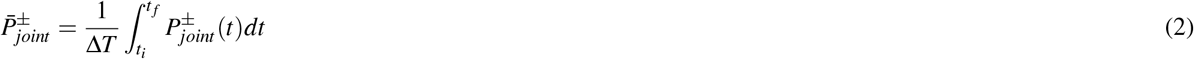

Two different temporal integration boundaries were used depending on the analysis objective. To assess segmental contributions over an identical analysis window, integration was performed over a full stride, defined from heel strike to the subsequent heel strike of the same leg (*t*_*i*_ = *t*_*HS*_, *t* _*f*_ = *t*_*nextHS*_). Stance and swing phases followed standard definitions. Integrated powers were normalized by stride duration (Δ*T* = *t*_*nextHS*_ – *t*_*HS*_).

For modeling EE estimation within the IMU framework, integration was restricted to the stance phase, from heel strike to toe off of the same leg (*t*_*i*_ = *t*_*HS*_, *t* _*f*_ = *t*_*TO*_). This choice reflects that IMU-based joint dynamics estimation is valid primarily during stance, where foot–ground contact mechanically constrains motion and reduces integration drift. Integrated powers were normalized by stance duration (Δ*T* = *t*_*TO*_ – *t*_*HS*_). Time-averaged positive, negative, and muscle efficiency-weighted joint powers were computed following the same procedure used in our previous work^16^, where *N*_*joint*_ denotes the number of joints in each segment. The same efficiency coefficients were also applied, with *η*^+^=4.00 for positive power and *η*^*−*^=-0.83 for negative power. These coefficients represent fixed constants derived from reported muscle efficiencies for positive and negative mechanical power, corresponding to efficiencies of 25% and 120%, respectively^27,28^.

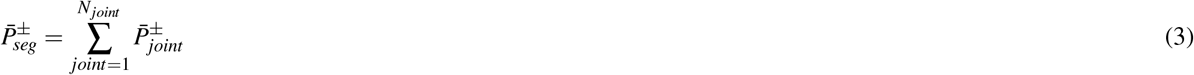

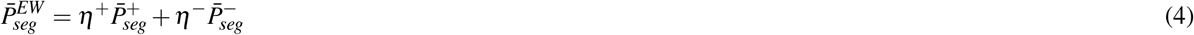

To convert efficiency-weighted power into estimated EE, a scale factor (*c*_*seg*_) was obtained for each segment by regressing efficiency-weighted power against measured EE. Leave-one-subject-out (LOSO) cross-validation was used to evaluate inter-subject consistency of the scale factors. In each LOSO iteration, the scale factor was obtained from the training subjects and applied to the held-out subject, resulting in subject-specific scale factors for evaluation. These scale factors were subsequently used in the IMU-based EE estimation framework. All computations were implemented in MATLAB R2022a (MathWorks, USA).

#### Machine learning to estimate EE from single IMU data

The proposed IMU-based EE estimation framework relies on a machine learning model that estimates stance-leg joint dynamics as an intermediate step. This model extends a previous network developed for estimating stance-leg dynamics during walking^29^ to accommodate running gait and to include swing-leg hip joint dynamics. The network inputs consisted of two-dimensional sagittal-plane accelerations from the sacrum-mounted IMU and the treadmill speed. As in prior work, treadmill speed was included to decouple biomechanical modeling from network complexity, and because running speed can be reliably obtained from wearable devices^30,31^. Since IMU-based estimation is valid only during stance phase, IMU signals were processed stride-by-stride, and the stance phase was extracted and interpolated to a normalized length. The neural network was implemented as a fully connected feed-forward artificial neural network (ANN) with a single hidden layer of 20 nodes. Inputs included treadmill speed (*v*) and normalized horizontal and vertical sacral displacements (*x, y* within [0, 1]). Outputs were stance-leg and swing-leg hip joint angles and torques. Displacements were computed by integrating IMU accelerations within each stance phase, and integration drift was removed by assuming equal sacrum height at heel strikes during steady-state running. Model training used a learning rate of 0.002, batch size of 128, and 50 epochs, and generalization performance was evaluated using LOSO cross-validation. Estimated joint angles and torques were filtered using a padded zero-lag 4th-order Butterworth filter (cutoff 10 Hz). Estimated joint power was computed as the product of filtered torque and numerically differentiated joint angle velocity. Efficiency-weighted power was computed using Equations 2.3 and 2.4, and subject-independent scale factors obtained from the training data were applied to obtain estimated EE (the hat symbol denotes estimated values).

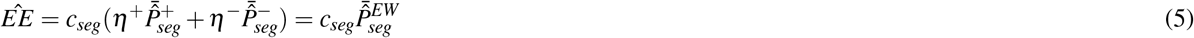

To compare the proposed biomechanically-informed framework with a direct data-driven approach, a comparison model that directly estimated EE from raw IMU signals was developed. Similar to the ANN architecture used in the proposed framework, the comparison model employed a single hidden layer feed-forward neural network, but without the intermediate biomechanical representation. Inputs included six IMU channels (triaxial acceleration and angular velocity), time, and treadmill speed; the output was a single scalar EE value (W/kg). Data preprocessing followed the same stance-phase extraction and interpolation used in the proposed framework. The comparison network consisted of a single hidden layer with 41 nodes, selected to match or exceed the representational capacity of the proposed intermediate biomechanics-based model while avoiding unnecessary overfitting given the dataset size. Training hyperparameters included a learning rate of 0.01, batch size of 16, and 50 training epochs. Training and evaluation used the same LOSO cross-validation procedure as the proposed framework.

## Results

As in walking, the stance leg, which is responsible for body weight support and propulsion, showed the largest contribution to whole-body joint mechanical power during running (Fig. 2*a*). During running, the stance leg, swing leg, and upper body contributed approximately 68.6%, 24.2%, and 7.2% of whole-body mechanical power, respectively. Compared with walking, the relative contribution of the stance leg decreased by approximately 16 percentage points, while the contributions of the swing leg and upper body increased by 12.3 and 3.7 percentage points, respectively (*p* < 0.05). More than 75% of the reduction in stance-leg power, for both positive and negative power, was offset by an increase in swing-leg power. Sagittal-plane power accounted for the majority of joint mechanical power in both walking and running (Fig. 2*b, c*). The contribution of sagittal-plane positive power did not differ significantly between the two gait modes (*p* > 0.05), and differences in other plane-wise components were small.

**Figure 2.**
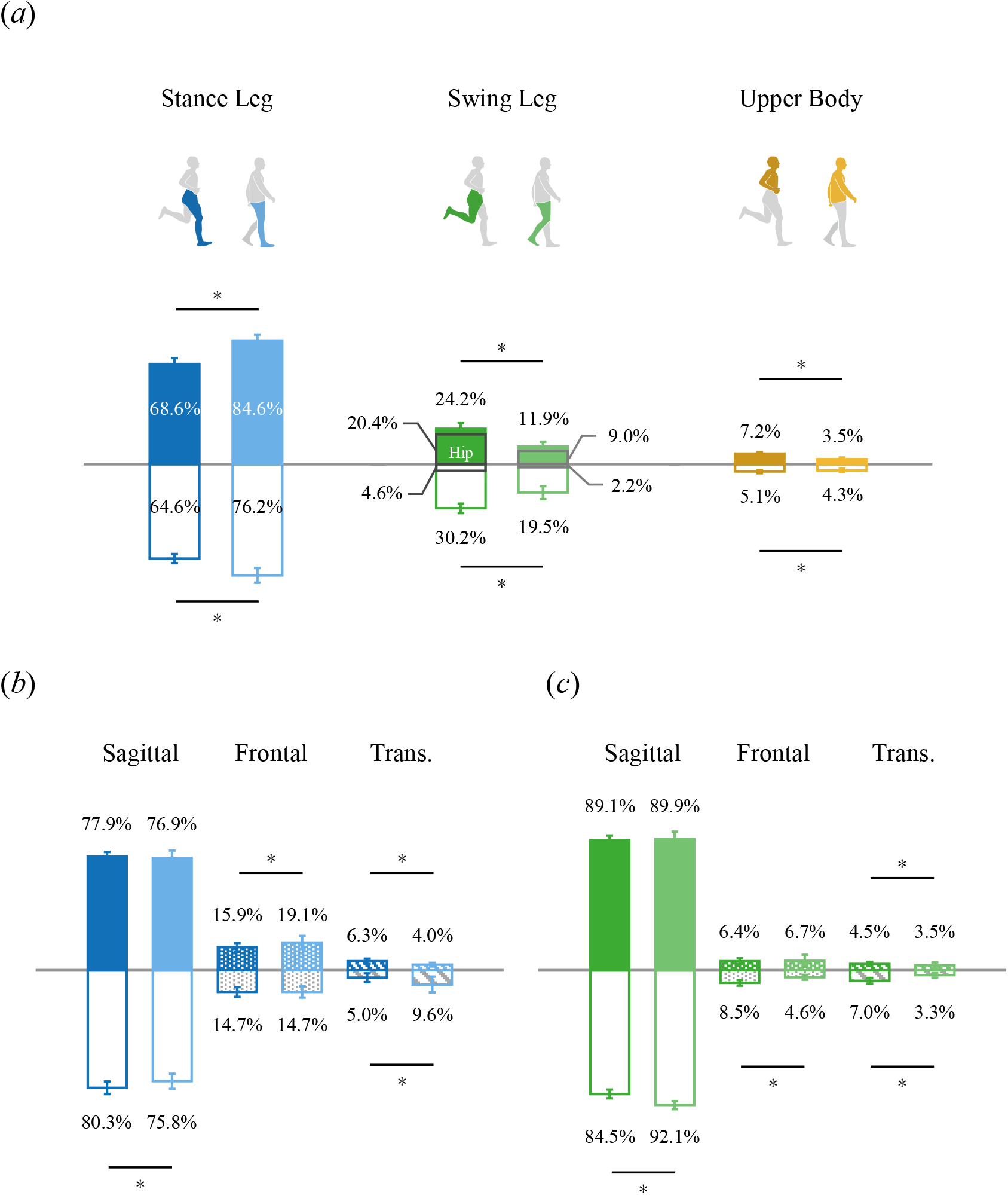
Segmental contributions to the whole-body joint mechanical power during a single stride of running (dark bars) and walking (light bars). (*a*) Percentage contributions of the stance leg (blue), swing leg (green), and upper body (yellow) to the time-averaged whole-body joint mechanical power. Percentage breakdown of (*b*) stance leg and (*c*) swing leg joint mechanical power into sagittal (filled), frontal (dotted), and transverse (dashed) plane components, relative to their respective total power. Shaded bars indicate positive power, and outlined bars indicate negative power. Error bars represent standard deviations across five gait speeds and eight subjects. Statistically significant differences (*p* < 0.05) are indicated with an asterisk (*).

The increased swing-leg power during running was primarily generated at the hip joint (Fig. 3*c*). The magnitude of hip joint power was markedly greater compared to walking and accounted for most of the increase in overall swing-leg power. Knee joint power also increased during running, with most of this increase occurring in negative power (Fig. 3*b*). In contrast, ankle joint power remained near zero in both walking and running, contributing minimally to swing-leg power (Fig. 3*a*).

**Figure 3.**
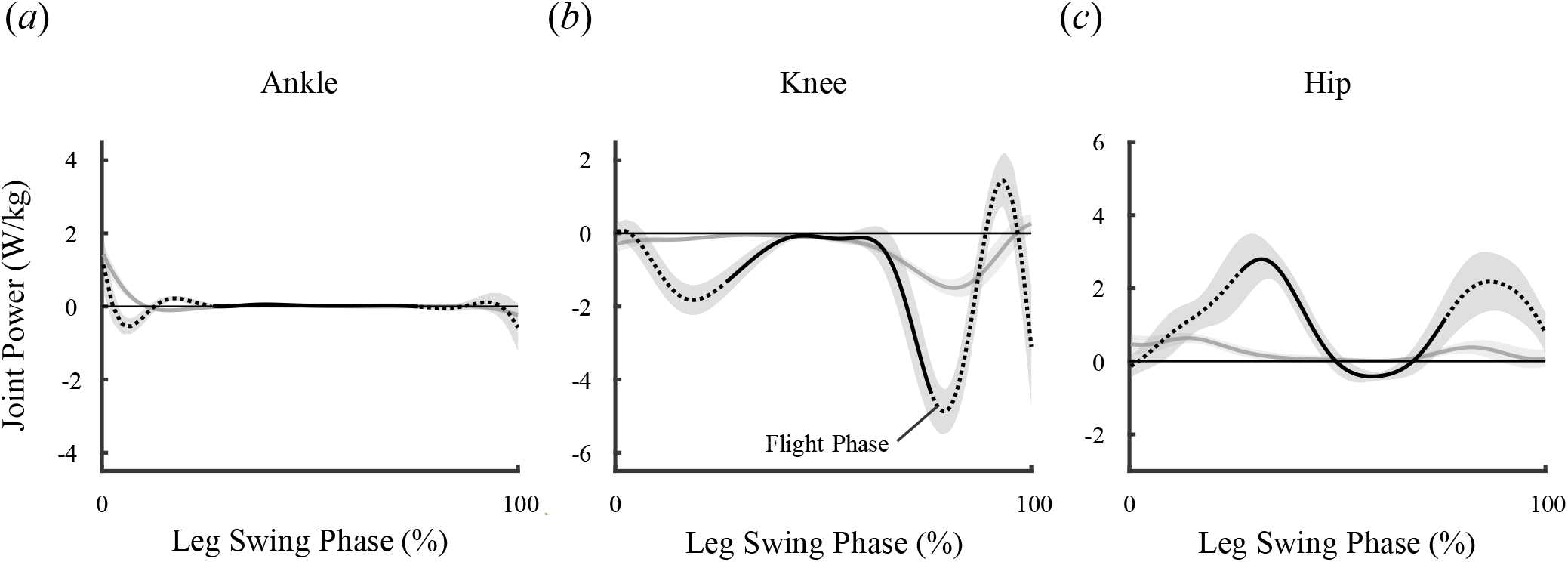
Mass normalized joint power of the swing leg at the (*a*) ankle, (*b*) knee, and (*c*) hip during the swing phase in running (black lines) and walking (grey lines). Solid lines and shaded areas indicate the mean and standard deviation across eight subjects during running at 3.1 m/s and walking at 1.5 m/s. Dashed lines indicate the flight phase within the full swing phase during running. Joint power is computed using the convention that positive joint angle and torque directions correspond to ankle plantar flexion, knee extension, and hip flexion.

Table 1 summarizes the scale factors (*c*_*seg*_), linear correlation coefficients (R), the significance of the correlations (*p*-values), and RMSE for each model. The scale factors showed very small intersubject standard deviations across all three models (*±* 0.003–0.007). The combination of stance leg and swing-leg hip joint model showed significantly lower RMSE than the stance leg only model. Figure 4 shows the relationship between measured EE and estimated EE based on joint mechanical power across all running speeds. High linear correlations with measured EE were observed for all three models: whole-body 3D power, the combination of stance-leg and swing-leg hip power (stance + swing hip), and stance-leg power alone (stance only). Subject-specific regression slopes also exhibited consistent patterns across models. The model using stance-leg and swing-leg hip power explained measured EE at a level comparable to the whole-body power model (R = 0.788 vs. R = 0.813), and stance-leg power alone also maintained a strong linear relationship with running EE (R > 0.75).

**Table 1.** Scale factors (*c*_*seg*_) for converting muscle efficiency–weighted joint mechanical power to measured EE, together with the linear correlations (R), the significance of the correlations (*p*-value), and estimation errors (RMSE, W/kg) between measured and estimated EE across all subjects and speeds. Estimated EE was computed as the product of the scale factor and the muscle efficiency–weighted joint mechanical power. Statistical significance of RMSE differences relative to the ‘stance leg only’ case is indicated with asterisks (\*\**p* < 0.01, \**p* < 0.05).

| Segment | $c_{seg}$ (s.d) | R | $p$-value | RMSE (s.d) |
| --- | --- | --- | --- | --- |
| Whole-body 3D | 0.252 (0.003) | 0.813 | $< 0.001$ | 1.252 (0.996) ** |
| Stance leg and swing-leg hip joint | 0.382 (0.005) | 0.788 | $< 0.001$ | 1.389 (0.986) * |
| Stance leg only | 0.435 (0.007) | 0.755 | $< 0.001$ | 1.553 (1.143) |

**Figure 4.**
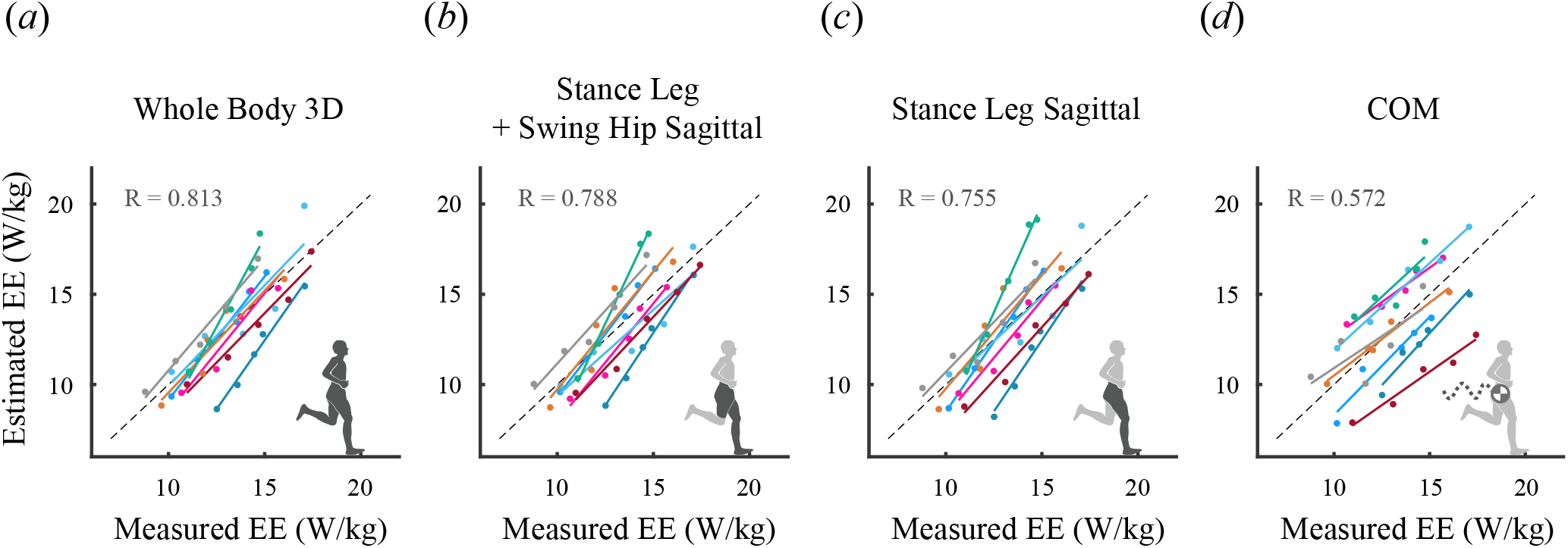
Comparison between measured EE via indirect calorimetry and estimated EE based on joint mechanical power for *(a)* the whole-body, (*b*) the sagittal stance leg + the swing-leg hip joint, (*c*) the sagittal stance leg alone, and (*d*) COM mechanical power. Colored circles represent individual subjects. Solid lines indicate subject specific linear regressions, and the dashed line represents the identity line where estimated EE equals measured EE.

Given that center of mass (COM) dynamics have traditionally served as a simple yet effective proxy for whole-body dynamics in gait^32,33^, it is important to evaluate whether COM power can capture EE variations across running speeds. Therefore, we also examined EE estimation based on COM power (Fig. 4*d*). COM power based estimates showed a markedly lower correlation (R = 0.572) and greater deviation from the identity line compared with joint power based estimates.

Using a lightweight ANN, sagittal-plane joint dynamics for the ankle, knee, and hip of the stance leg, as well as the hip of the swing leg, were accurately estimated from a single sacrum-mounted IMU. Compared to ground truth data obtained from inverse dynamics analysis of motion capture data, the mean RMSEs for joint angle, torque, and power were 4.43, 4.88, 6.47 °, 0.31, 0.31, 0.29 Nm/kg, and 2.30, 1.82, 1.27 W/kg, respectively (Table 2). In addition, the errors of newly estimated swing-leg hip joint dynamics were 6.96 ° for angle, 0.15 Nm/kg for torque, and 0.65 W/kg for power.

**Table 2.** Estimation errors of the stance-leg sagittal joint and swing-leg sagittal hip joint dynamics obtained using an ANN and a single sacrum-mounted IMU. Mean RMSE values (with standard deviations in parentheses) across all subjects are presented for each running speed and for each joint (stance ankle, knee, hip, and swing hip), and averaged value across all running speeds.

| Variable | Speed (m/s) | Stance Ankle | Stance Knee | Stance Hip | Swing Hip |
| --- | --- | --- | --- | --- | --- |
| Angle (°) | 2.5 | 4.68 (2.08) | 5.49 (2.27) | 6.08 (2.62) | 6.53 (2.71) |
|  | 2.8 | 3.97 (1.39) | 5.16 (2.89) | 5.84 (2.62) | 5.94 (3.16) |
|  | 3.1 | 4.09 (1.50) | 4.07 (2.38) | 7.60 (4.83) | 7.88 (5.51) |
|  | 3.4 | 4.40 (1.63) | 4.05 (2.20) | 6.90 (3.74) | 7.88 (4.53) |
|  | 3.7 | 5.01 (1.33) | 5.65 (3.30) | 5.94 (2.53) | 6.55 (3.19) |
|  | Average | 4.43 (1.65) | 4.88 (2.73) | 6.47 (3.46) | 6.96 (4.04) |
| Torque (Nm/kg) | 2.5 | 0.31 (0.15) | 0.32 (0.23) | 0.30 (0.18) | 0.15 (0.04) |
|  | 2.8 | 0.24 (0.09) | 0.28 (0.14) | 0.24 (0.09) | 0.14 (0.04) |
|  | 3.1 | 0.30 (0.11) | 0.29 (0.10) | 0.31 (0.17) | 0.15 (0.08) |
|  | 3.4 | 0.34 (0.13) | 0.32 (0.12) | 0.34 (0.18) | 0.14 (0.06) |
|  | 3.7 | 0.35 (0.09) | 0.34 (0.18) | 0.27 (0.10) | 0.14 (0.04) |
|  | Average | 0.31 (0.12) | 0.31 (0.16) | 0.29 (0.15) | 0.15 (0.05) |
| Power (W/kg) | 2.5 | 1.95 (1.15) | 1.58 (0.60) | 1.20 (0.96) | 0.63 (0.24) |
|  | 2.8 | 1.75 (0.59) | 1.61 (0.57) | 0.89 (0.39) | 0.57 (0.23) |
|  | 3.1 | 2.11 (0.71) | 1.80 (0.51) | 1.34 (0.75) | 0.54 (0.24) |
|  | 3.4 | 2.66 (0.84) | 1.92 (0.68) | 1.61 (0.89) | 0.63 (0.26) |
|  | 3.7 | 3.02 (0.91) | 2.20 (0.82) | 1.33 (0.61) | 0.88 (0.29) |
|  | Average | 2.30 (0.98) | 1.82 (0.68) | 1.27 (0.78) | 0.65 (0.28) |

EE estimation based on joint mechanical power calculated from a single sacrum-mounted IMU showed high linearity (R = 0.838), with estimated values closely aligned with the identity line (Fig. 5*a*). The RMSE was 1.40 W/kg, and residuals exhibited no significant correlation with measured EE (*p* > 0.05). In contrast, the direct data-driven method (*b*), which estimates EE directly from IMU signals without incorporating lower-limb joint dynamics, showed a lower correlation coefficient (R = 0.717) with an RMSE of 1.32 W/kg (Fig. 5*b*). The difference in RMSE between two methods was not statistically significant (*p* > 0.05). Estimation variability of the direct data-driven method increased at higher speeds, and the residuals displayed a significant linear relationship with measured EE (*p* < 0.01).

**Figure 5.**
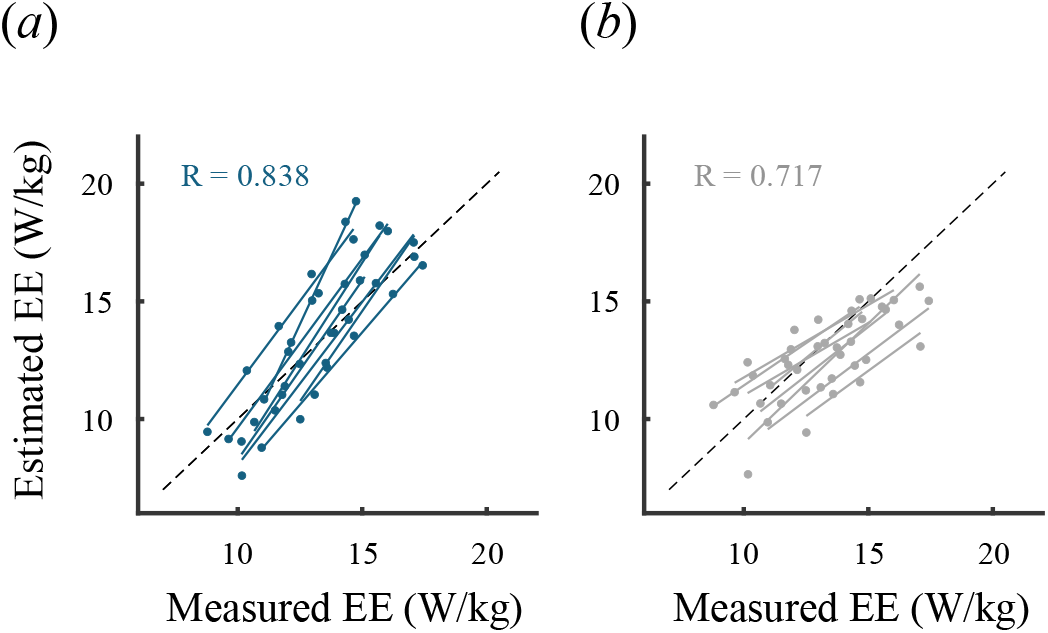
Comparison between measured EE via indirect calorimetry and estimated EE based on joint mechanical power computed using a single sacrum mounted IMU. (*a*) Estimated EE calculated from stance-leg sagittal plane joint power (blue). *(b)* Estimated EE obtained directly from IMU signals using a data-driven machine learning model (grey). Circles represent individual subjects. Solid lines indicate subject specific linear regressions, and the dashed line represents the identity line where estimated EE equals measured EE.

## Discussions

Although the stance leg remained the dominant contributor to mechanical power during running, the ratio of stance-leg power to whole-body power was reduced, accompanied by a significant increase in the contribution of the swing leg (Fig. 2*a*). This shift highlights both the shared reliance on stance-leg dynamics and the greater involvement of swing-leg dynamics that characterizes running compared with walking. Because the stance phase in running occupies a substantially smaller proportion of the stride^34^ and occurs at higher forward speed, greater collision losses arise^35^, and the corresponding energy must be rapidly restored to maintain forward progression. These demands explain the high mechanical power of the stance leg, which remains the dominant contributor in running^36^. This pattern is consistent with previous electromyography findings showing increased activation of major lower-limb muscles, including the gluteal muscles, during running compared with walking^37^. In walking, the swing leg relies on pendular dynamics enabled by the sufficiently long stance phase, resulting in relatively low muscle activation demands^38^. In contrast, running requires larger joint excursions and higher muscle activation not only for rapid stride generation but also for flight propulsion and landing preparation^37,39,40^. These neuromuscular characteristics are consistent with the increased swing-leg mechanical power observed in this study (walking ∼12% *→* running ∼20–24%). Furthermore, during the flight phase of running, both legs function as swing legs, which structurally increases the relative contribution of the swing leg over the entire stride.

Approximately 70% of the increase in upper-body mechanical power during running originated from the torso. The torso undergoes slight flexion during early-stance collision and re-extension during propulsion, generating substantial mechanical power due to its large mass. Despite the distinct segment-level differences between walking and running, plane-wise contributions were largely similar across both conditions, with sagittal-plane power remaining dominant (Fig. 2*b, c*). Taken together, these findings highlight the central role of the stance leg in mechanical power-based EE estimation and suggest that incorporating swing-leg power, particularly during running, may further improve the representation of whole-body mechanical power.

The large hip joint power observed during running (Fig. 3*c*) likely reflects the increased demand for rapid limb acceleration and deceleration of leg swinging. The hip joint produced high positive power in the early swing phase to accelerate the lower limb forward, followed by negative power in the late swing phase to decelerate the limb in preparation for landing. During running, the hip flexors and extensors of the swing leg must contract more rapidly and forcefully to generate stride length and prepare for the subsequent stance phase^41^, with particularly high hip flexion power required to overcome inertial forces during the flight phase. The magnitude of hip joint power was markedly greater compared to walking and accounted for most of the increase in overall swing-leg power. These demands are consistent with previous studies reporting elevated activation of muscles surrounding the hip during running^37,40,42^. The large negative power of knee joint observed during mid flight phase (Fig. 3*b*) serves to decelerate the extending shank during the swing phase^43^. However, because negative power is weighted with a substantially smaller efficiency coefficient than positive power (*η*^*−*^ = -0.83 vs. *η*^+^ = 4.00^27,28^), the increase in knee joint negative power contributes relatively little to the efficiency-weighted joint power used for EE estimation. These results indicate that the increased mechanical contribution of the swing leg during running is primarily attributable to hip joint power, supporting the rationale for additionally considering swing-leg hip power alongside stance-leg power in mechanical power based EE estimation.

A scale factor less than 1 indicates that the product of joint mechanical power and muscle efficiency exceeds the actual metabolic cost measured via respiratory gas analysis. In other words, the estimates based on mechanical power and muscle efficiency calculated in this study tend to overestimate actual metabolic cost by approximately fourfold when using whole-body mechanical power and about twofold when using stance-leg mechanical power. This overestimation can be interpreted from two perspectives. First, overestimation in the mechanical power calculation itself, and second, overestimation of the proportion of mechanical power attributable to active muscle contraction. First, the mechanical power calculation in this study summed the time-averaged joint power computed separately for each joint, and therefore did not account for energy transfer between joints via biarticular muscles. This would lead to overestimation of efficiency-weighted power. Previous research reported that the difference between maximum and minimum assumptions for inter joint energy transfer is approximately 20%^14^. Applying a correction for average energy transfer effects, the scale factor based on stance-leg mechanical power is estimated to be approximately 0.5.

Second, even with this correction, a scale factor of approximately 0.5 suggests that the mechanical power attributable to active muscle contraction is of a comparably similar magnitude. This implies that mechanical power during running is generated through a mixed mechanism involving both muscle activation and elastic recoil, with their relative contributions appearing to be of comparable scale. Numerous studies have reported substantial contributions of tendon elastic strain energy during running^18,19,44–46^. In particular, Riddick and Kuo^14^ quantitatively evaluated the contribution of mechanical power to EE during running and set the contribution of elastic recoil at a provisional nominal value of 50%, which is consistent with the range observed in the present study. Furthermore, elastic recoil is known to play a much smaller role during walking than running^45,47^, and the scale factor for walking has been reported to be approximately twice that of running^16^, supporting the present interpretation based on elastic energy contribution.

A limitation of this approach is that metabolic energy expenditure components not captured by mechanical power may influence the scale factor. For example, if factors such as force rate cost are included^14,15^, the scale factor values reported here may change. In related sense, the interpretation of comparable contributions from active muscle work and elastic recoil may be modestly influenced by metabolic processes that do not produce external work (e.g., isometric force production, co-contraction). Additionally, as noted in the methods, the analysis in this study focused on using stance-phase information, with IMU-based EE estimation in mind, and therefore only stance-phase mechanical power was used in calculating the scale factors rather than the entire stride. Nevertheless, as shown in Figure 4, mechanical power during the stance phase accounts for the dominant proportion of total mechanical power, so the difference between using stance-phase power and full stride power is expected to be small.

The high linear correlation with measured EE and the consistent pattern of subject-specific regression slopes shown in figure 4 indicate that the primary determinants of running EE are concentrated in lower-limb segmental dynamics. Because propulsion forces during running are generated primarily during the stance phase, stance-leg power contains sufficient information to represent whole-body EE. This is supported by the high linearity (R > 0.75) and the approximately 70% mechanical contribution discussed earlier as well as previous studies^16,48^. Including swing-leg hip power produces a significant improvement in estimation performance (*p* < 0.05), suggesting that active torque production by the hip flexors and extensors during the swing phase imposes a non-negligible metabolic demand. This observation aligns with previous study identifying swing-leg hip power as a meaningful component of running cost^49^. In particular, the rapid torque-rate changes at the hip and the energetic demands of swing acceleration and deceleration are known contributors to metabolic cost^50^, and the present findings provide quantitative support for this mechanism. Although not shown in the presented results, a model incorporating the full swing leg (ankle, knee, and hip) yielded R and RMSE values nearly identical to those of model B, with no statistically significant differences.

As discussed earlier, the ankle contributes minimally to swing motion, and the knee is dominated by negative power, whose efficiency weight is roughly one-fifth that of positive power^27^, resulting in a limited influence on EE. Thus, the performance improvement associated with adding swing-leg information is driven primarily by the hip joint, which generates predominantly positive power, and the essential contribution of the swing leg appears to be fully captured by the hip joint alone.

These findings highlight the importance of joint-level dynamics in explaining running EE. In contrast, COM dynamics, often used as a simple proxy for whole-body dynamics in gait, showed limited ability to capture EE, as reflected by the poor estimation performance (Fig. 4*d*). To understand this limitation, we examined the relationship between COM power and whole-body joint power across speeds. While lower-limb joint power consistently accounted for a large proportion (about 70%) of whole-body power, COM power accounted for a much smaller proportion and gradually decreased from about 30% as running speed increased (Fig. 6). These findings indicate that the rise in metabolic cost with running speed is driven primarily by active mechanical power at the lower-limb joints, whereas COM dynamics play a comparatively limited role.

**Figure 6.**
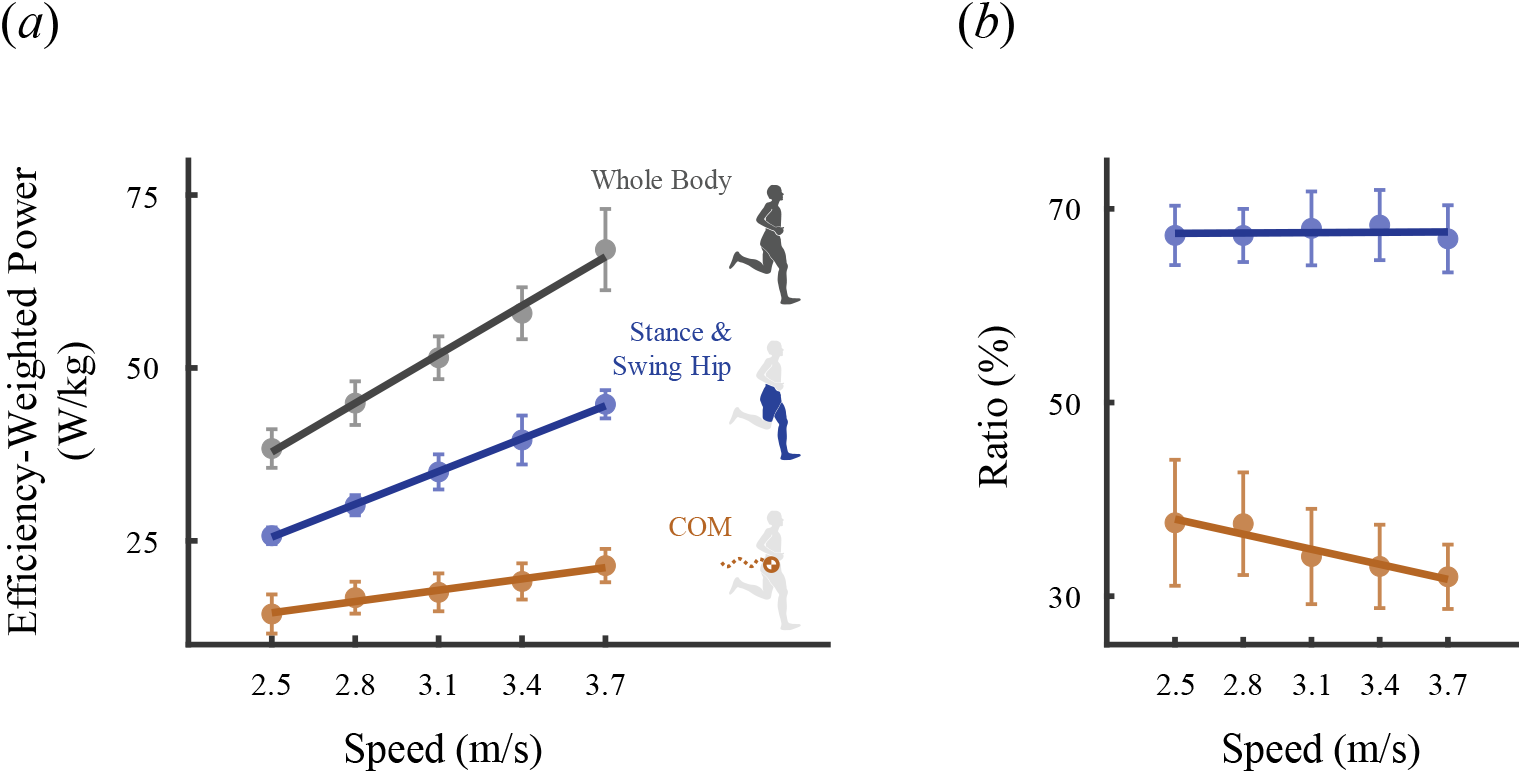
(*a*) Muscle efficiency-weighted power at different running speeds for the whole-body (black), stance leg combined with the swing-leg hip joint (blue), and the center of mass (COM, orange). (*b*) Ratios of the limb joint power (blue in *a*) and COM power (orange in *a*) to the whole-body power (black in *a*) across running speeds. Error bars in (*a*) and (*b*) indicate standard deviations across eight subjects.

COM-based mechanical models are useful for explaining overall energy transformation by simplifying the body as a single point mass. However, they cannot capture limb-specific movements or the mechanical power generated at individual joints. During running, active mechanical power at the lower-limb joints increases substantially due to rapid limb accelerations and short ground contact times^34,46,51^, and this joint-level power is the primary driver of the speed dependent rise in EE. In contrast, COM power primarily reflects vertical oscillation and small fluctuations in anteroposterior velocity of the body’s center of mass, and therefore shows minimal change with increasing speed during steady-state running at constant velocity. These differences were directly reflected in EE estimation performance, which showed poor sensitivity to speed-induced changes in EE. By contrast, estimates derived from the combination of stance-leg and swing-leg hip joint power closely tracked measured EE across the full range of running speeds, confirming that joint-level dynamics contain the essential mechanical information required for accurate EE prediction. Overall, these results demonstrate that COM-based dynamics are insufficient for modeling metabolic cost during running, where limb-level mechanical power is the primary driver of energetic demand. For single IMU applications, this supports the validity of the proposed stance-leg and swing-leg hip power based approach, which more accurately captures the mechanical determinants of EE and explains speed dependent metabolic changes that COM-based methods cannot represent.

The estimation accuracies of the lightweight ANN shown in Table 2 are comparable to prior IMU-based studies that estimated specific joint dynamics during running^52–55^. Although the network was originally designed for walking^29^, it was successfully extended to running conditions without substantial performance degradation. In the present study, swing-leg hip joint dynamics were newly estimated, which were not included in the original walking-based framework^16,29^. Swing-leg hip joint power showed pronounced subject-specific differences during early stance (not shown), and a substantial portion of the total estimation error originated in this interval. Because the swing leg is dynamically coupled to the center of mass, a certain level of estimation was still possible from a single sacrum-mounted IMU. However, a deeper biomechanical representation of swing-leg dynamics is expected to directly contribute to improvements in estimation accuracy.

The high linearity, accuracy, and residual characteristics observed in the estimation results indicate that the biomechanically-informed method achieves stable performance across walking and all running speeds without systematic overestimation or underestimation. The RMSE of 1.40 W/kg is also competitive with values reported in previous multi sensor or heart rate based running EE estimation studies^22,24,25^. The direct data-driven method produced similar or slightly lower RMSE, but showed lower sensitivity to the subject-specific increase in EE with speed. This behavior can be explained by a mean regression effect, in which predictions tend to be pulled toward the central range of the data when the model learns solely through label-based error minimization. Estimation variability increased at higher speeds, and the residuals displayed a significant linear relationship with measured EE (*p* < 0.01), indicating a proportional bias in which higher EE values were systematically underestimated. These findings demonstrate that EE during running can be practically estimated using only a single sacrum-mounted IMU. As in walking, when data are limited and a simple machine learning model is used, incorporating lower-limb joint dynamics provides higher accuracy and better generalization than direct data-driven approaches.

## Conclusions

We confirmed that lower-limb mechanical power is strongly correlated with EE during running and quantified the relative contributions of segments. This relationship was quantitatively characterized through the scale factor linking efficiency-weighted power to EE, supporting previous observations^14,15^ that the metabolic cost associated with active muscle work remains substantial despite the contribution of elastic recoil. Furthermore, we quantitatively demonstrated that the stance leg contributes dominantly to EE across a wide range of speeds, extending from walking^16^ to running, while swing-leg hip joint provides additional explanatory information that improves estimation performance. These findings simultaneously reflect both the shared and distinct characteristics of walking and running in terms of dynamics and energetics. Moreover, they confirm that the biomechanically-informed framework previously demonstrated for walking can be extended to running, supporting the general applicability of this approach across different gait modes. Beyond these biomechanical insights, the present results also demonstrate the practical feasibility of estimating EE during running from a single sacrum-mounted IMU. In this regard, this study provides a foundation for the practical implementation of IMU-based metabolic monitoring during running, with potential applications in sports science, human activity monitoring, and rehabilitation.

## Ethics

Each participant gave written informed consent as approved by the KAIST Institutional Review Board (KH2023-250).

## Data availability

Experimental data and code for analysis are available from the Zenodo digital repository^56^.

## Declaration of AI

We have not used AI-assisted technologies in creating this article.

## Author contributions statement

J.J. contributed to the data collection, processing, design of the analysis, performed the analysis, and wrote the manuscript.

H.L. contributed to the machine learning analysis. S.P. designed the overall study and analysis methods and organized the manuscript. All authors reviewed and approved the final manuscript.

## Additional information Competing interests

The author(s) declare no competing interests.

